# Gene expression is a poor predictor of the metabolite abundance in cancer cells

**DOI:** 10.1101/2021.12.19.473333

**Authors:** Huaping Li, Jayne A. Barbour, Xiaoqiang Zhu, Jason W. H. Wong

## Abstract

Metabolic reprogramming is a hallmark of cancer characterized by global changes in metabolite levels. However, compared with the study of gene expression, profiling of metabolites in cancer samples remains relatively understudied. We obtained metabolomic profiling and gene expression data from 454 human solid cancer cell lines across 24 cancer types from the Cancer Cell Line Encyclopedia (CCLE) database, to evaluate the feasibility of inferring metabolite levels from gene expression data. For each metabolite, we trained multivariable *LASSO* regression models to identify gene sets that are most predictive of the level of each metabolite profiled. Even when accounting for cell culture conditions or cell lineage in the model, few metabolites could be accurately predicted. In some cases, the inclusion of the upstream and downstream metabolites improved prediction accuracy, suggesting that gene expression is a poor predictor of steady-state metabolite levels. Our analysis uncovered a single robust relationship between the expression of nicotinamide N-methyltransferase (*NNMT*) and 1-methylnicotinamide (MNA), however, this relationship could only be validated in cancer samples with high purity, as *NNMT* is not expressed in immune cells. Together, our findings reveal the challenge of inferring metabolite levels from metabolic enzyme levels and suggest that direct metabolomic profiling is necessary to study metabolism in cancer.

## Introduction

Metabolism refers to all biochemical reactions taking place in live cells and organisms. Metabolites are the by-products or products of cellular metabolic activities, including nucleotides, amino acids, lipids, saccharides and other small molecules. Cancer cells tend to modulate their metabolism to meet the high anabolic requirements of rapid proliferation^1,2^. The Warburg effect describes the excessive intake of glucose and its conversion to lactate and has been observed in cancer for almost a century^3^. This metabolic program promotes cancer progression^4,5^. The cancer metabolic properties are impacted by intrinsic and extrinsic factors, such as nutrient and waste concentration, pH and oxygen tension. Alterations in metabolic pathway activities, also called metabolic reprogramming, has been reported as a hallmark of cancer^6^. The metabolic interactions *in vivo* work symbiotically to support cancer cells’ growth and maintenance, and even contribute to immune evasion^7^. Metabolic activities of pathways such as glutathione metabolism^8^, and the well-studied glucose-alanine cycle^9^ can partly account for cancer progression. These highlight that metabolite detection is important for disease diagnosis, cancer typing, as well as drug development.

Over the past decade, numerous genomes, epigenomes, and proteomes of large cancer cohorts have been profiled, with many malignancy-related hallmark molecular alterations reported across various cancer types. Metabolism plays a vital role in cancer development, however, there have been no pan-cancer metabolomics studies that match the research efforts of large scale projects such as The Cancer Genome Atlas (TCGA)^10,11^ and Clinical Proteomic Tumor Analysis Consortium (CPTAC)^12^. Major advances have been made in analytical techniques, such as liquid chromatograph-mass spectrometry (LC–MS), gas chromatography (GC)–MS, and LC–nuclear magnetic resonance (NMR), yet precise metabolite profiling remains challenging. This may be partly due to the heterogeneity of metabolites present in cells, resulting in a lack of standardization in metabolomic profiling. Thus, relatively few metabolomics studies have been performed in large cohorts of cancer samples.

Although several computational tools have been developed for metabolite prediction^13,14^, their primary function is focused on predicting biotransformation of exogenous metabolites, rather than predicting the abundance of endogenous metabolites in cells. A more recent study found relationships between transcriptional regulator activity and metabolite levels, however, the associations were limited to 53 cancer cell lines^15^. On the other hand, numerous studies on cancer metabolism have solely used gene and protein expression to infer metabolic activity without direct profiling of metabolite levels^16,17,18,19^. However, gene expression patterns may not intuitively reflect the activity of metabolic pathways^20^, where the reaction rate or environmental factors may also have a substantial impact. Another issue in previous cancer metabolism studies is that most research was focused on a specific cancer type^21,22,23,24^ or metabolic pathway^25,26^, limiting a comprehensive understanding of metabolism reprogramming.

Cancer cell lines at some level reflect cancer processes *in vitro* within a relatively regulated environment. As a previous study has shown, the nutrient milieu from cell-cultured media can reduce the impact of cell-intrinsic metabolic preferences^27^. Recently, larger metabolomic data from cancer cell lines has been collected^28^. The Cancer Cell Line Encyclopedia (CCLE) Database (https://sites.broadinstitute.org/ccle/) stores multi-omics data for over 1000 cell lines and is a valuable resource to learn the interactions between gene expression patterns and metabolism. In this study, using data from the CCLE, we rigorously investigated whether metabolites can be modelled by gene expression data, with the aim to predict metabolite levels in datasets without direct acquisition of metabolomics data.

## Results

### Metabolomic profiles are relatively stable across adherent cancer cell lines

After sample filtering and data pre-processing (see Materials and Methods), 454 adherent-growth solid cancer human cell lines (Fig 1a) with 225 metabolites and 13,403 coding genes expression profiles were used in this study (Supplementary Table 1). Hierarchical clustering analyses show no cancer type-specific metabolism pattern (Fig. 1b). We then assessed the variability and relationship between metabolites by calculating the coefficient of variation (CV) across all 454 cell lines. Except for 9 metabolites, most have low CV (< 0.1) (Fig. 1c, Supplementary Table 2), which suggests that most metabolites were stable across pan-solid cancer cell lines. Pearson correlation analyses were also performed for every pair of metabolites (R = -0.598 to 1, mean = 0.0018; Supplementary Fig. 1). Lipids related to triacylglycerol and ceramide were closer to each other, as well as some amino acids. However, except for some lipids, most of the 225 metabolites were relatively independent and showed weak correlations (absolute value of Pearson score < 0.5) with other metabolites.

**Figure 1.**
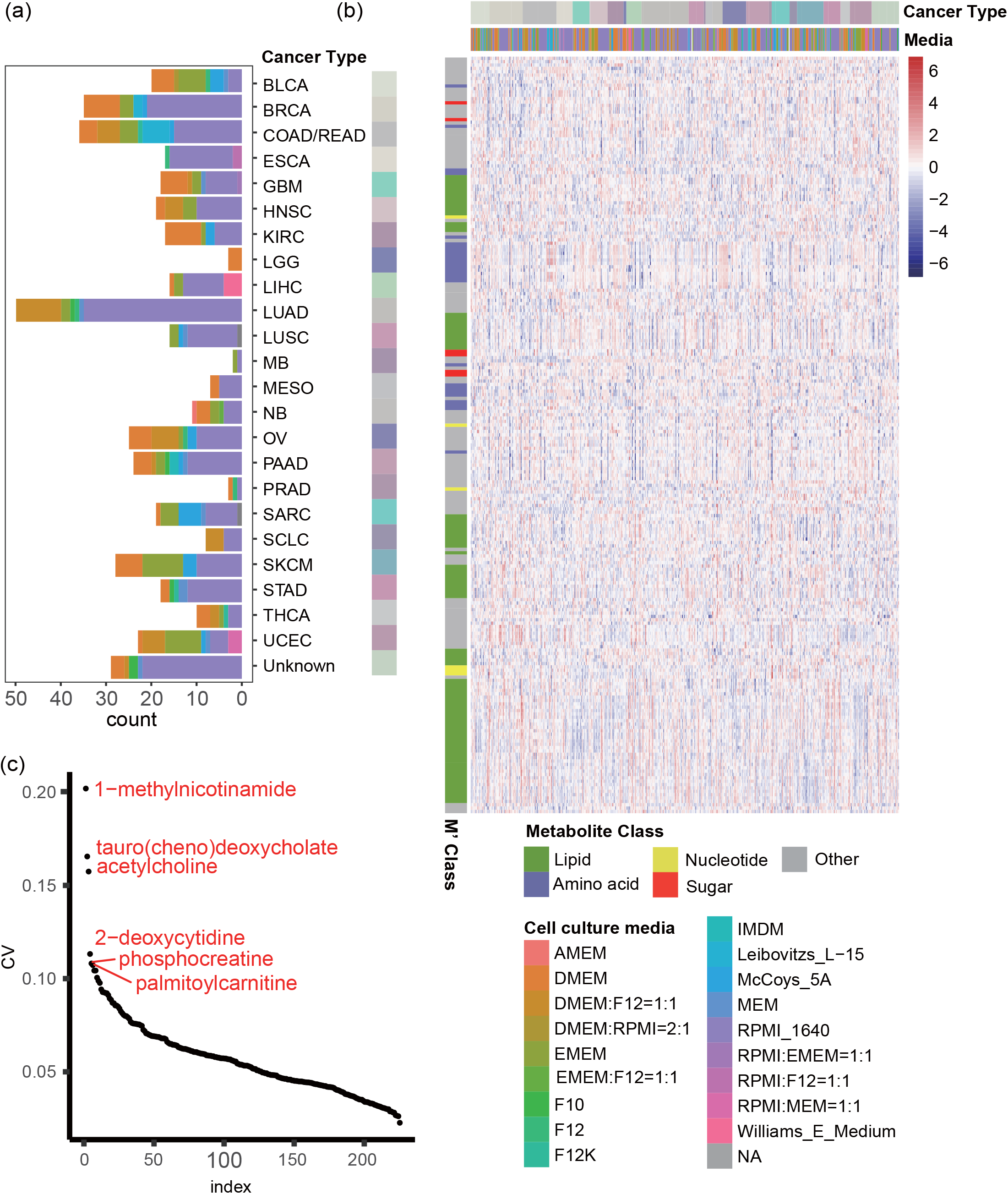
statistical analysis based on CCLE metabolomic profiles. 454 Cell lines (over 24 solid cancer types, 18 media types) were utilized in this study. (a) Cancer types distribution among adherent cultured cell lines. (b) Heatmap plot from cancer cell lines metabolomic profiling. Rows are metabolites, columns are cell lines grouped by cancer types, no cancer type-specific metabolism pattern is shown. (c) Ranked metabolites based on the estimated coefficients of variance (CV). The “Cancer Type” were labelled based on TCGA Study Abbreviations.

### Cell culture media does not profoundly influence metabolomic profiles

Up to 18 common media were used to maintain the solid cancer cell lines in culture by the CCLE (Supplementary Table 3). *RPMI 1640*(49.7%) is the major media used, followed by *DMEM* (15.0%) and *EMEM* (11.7%) (Fig 1a). We first investigated whether the media components and clinical traits will influence the metabolite levels. In total, 55 components from media were analyzed, along with clinical information including inferred ethnicity, patient age, pathology, cancer type, mutation rates and doubling time (hr). Meanwhile, we evaluated the variance inflation factor (VIF) among those features. Most features generated large VIF values (e.g. D-Calcium pantothenate VIF = 670 and Folic Acid VIF = 46), this indicated a high degree of multicollinearity among features. Therefore, using media components and clinical traits data, we trained a step-wise covariate linear regression model against each metabolite (Fig. 2b). R-squared (R^2^) was used to evaluate the proportion that the metabolite levels would be influenced by the features (Supplementary Table 4). Seven metabolites had relatively higher R^2^ (R^2^ >0.5). However, 91.6% (206 of 225) predicted metabolites had an R^2^ lower than 0.3, indicating that they were not greatly influenced by cell culture media or clinical traits.

**Figure 2.**
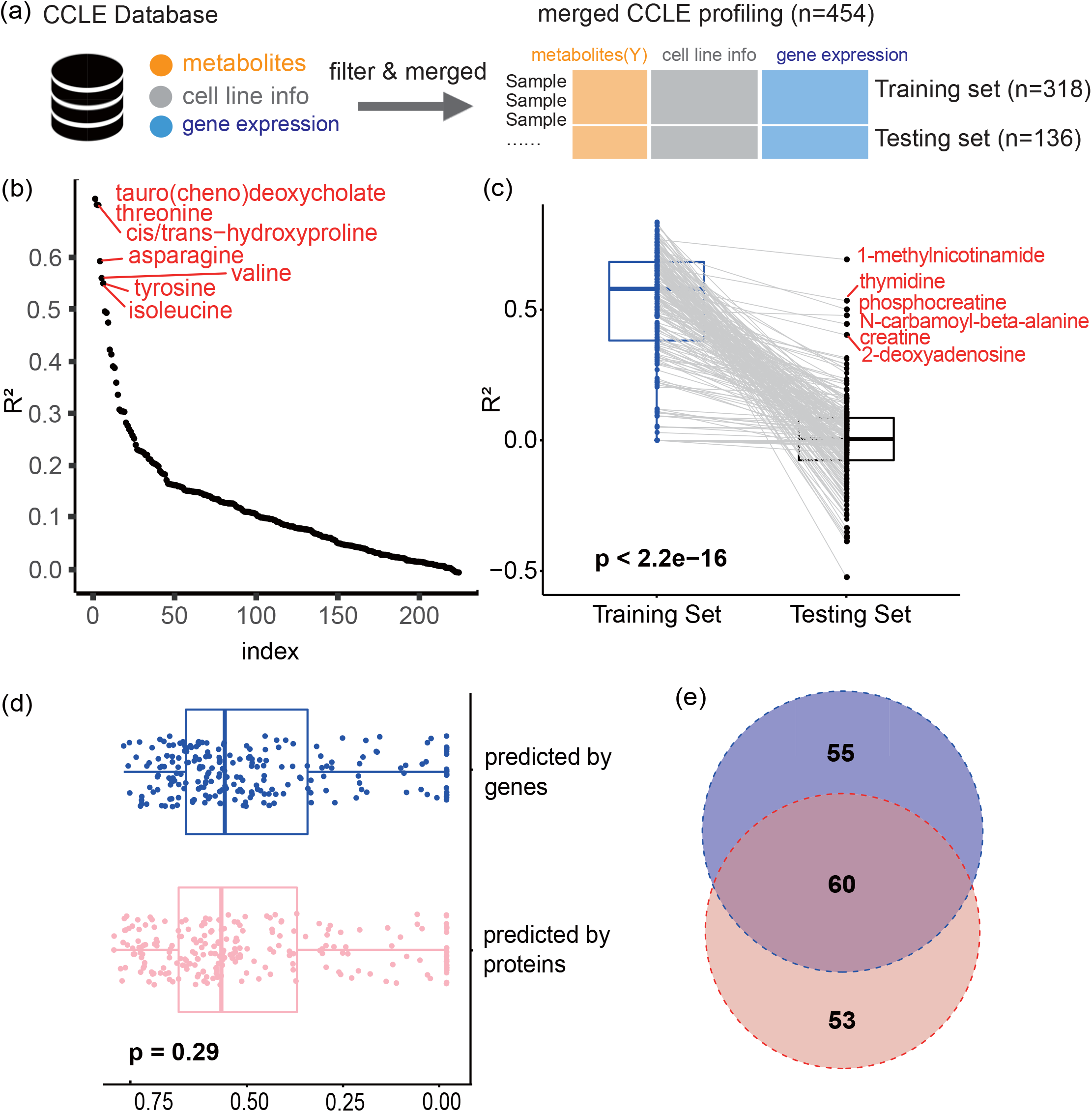
covariates analysis and global gene-based model training result. a) 454 human solid cancer cell lines were randomly assigned into training set (n=318) and testing set (n=136) after sample filtering and data merging; b) R^2^ distribution of step-wise linear regression models, only 7 metabolites have R^2^ higher than 0.5. c) R^2^ distribution of global gene-based Lasso regression prediction models, the median of R^2^ in training set and testing set were 0.58 and 0, respectively. d) R^2^ distribution of global gene-based Lasso models (blue) and proteomics-based Lasso models (red). e) Venn diagram of common predictable metabolites (Top 50% high R^2^ metabolites in testing set) between global gene-based (blue) and proteomics-based (red) Lasso models.

### Few metabolites can be predicted by gene expression levels

Since culture media components and clinical traits were not found to significantly affect the metabolite levels, we proceeded to develop gene-based models without additional covariates. The 454 cell lines were randomly split into training (n=318) and testing set (n=136). Feature selection was first performed by univariable linear regression analysis (see Materials and Methods). For each metabolite, we ranked the genes from high to low R^2^ and selected the top 10% genes only. Using the selected genes, we implemented a global gene-based *LASSO* regression model to predict the targeted metabolite level, and then validated it in the testing set. As the number of selected genes was different from every metabolite, the size of trained features in *LASSO* model changed.

We regarded R^2^ as the model evaluation criteria. The R^2^ from the training set ranged from 0 to 0.85, with a median value of 0.58, while the R^2^ from the testing set ranged from -0.524 to 0.692, with a median of 0 (Fig. 2c; Supplementary Table 5). Only 6 metabolite models stood out (R^2^ score >0.4) in the testing set. These 6 metabolites were not influenced by cell culture conditions as they had low R^2^ in step-wise linear regression models (Table 1). However, 51% (115 of 225) metabolites generated R^2^ equal to or lower than 0 in the testing set. In other words, more than half of the profiled metabolites were unpredictable by gene expression data.

**Table 1.** Metabolites with high R-squared(R^2^) from the prediction models.

|  | **Global gene-based LASSO model** |  | **covariates**<br>**linear model<sup>^</sup>** |
| --- | --- | --- | --- |
| **Metabolite** | **TrainingSet_ $R^2$** | **TestingSet_ $R^2$** | **$R^2$** |
| **1-methylnicotinamide** | 0.769 | 0.692 | 0.0382 |
| **thymidine** | 0.666 | 0.534 | 0.0408 |
| **phosphocreatine** | 0.728 | 0.501 | 0.0027 |
| **N-carbamoyl-beta-alanine** | 0.497 | 0.478 | 0.0449 |
| **creatine** | 0.592 | 0.445 | 0.0980 |
| **2-deoxyadenosine** | 0.581 | 0.403 | 0.0222 |

<sup>^</sup>Covariates linear model shows the impact from cell culture condition and clinical traits to metabolite levels.

Despite the R^2^ of global gene-based *LASSO* prediction models being typically low in the training set, many are still significantly higher than random expectation (permutation tests, all R^2^ equal to 0). This implied the gene regulation still contributes to metabolite level prediction but is not a major factor in most cases. In addition to feature selection, we also trained multivariable *LASSO* prediction models based on all 13k profiled genes (Supplementary Fig. 2A). By comparing the R^2^ histogram, it is clear that the overall performance of prediction models is better than those without a feature selection process.

To test whether protein levels will be a better predictor than gene expression, the same approaches were applied using proteomic data. The CCLE also generated mass spectrometry-based proteomes of 378 cell lines^29^, with 351 in common with cell lines with the metabolomic profiles. After removing the lowly correlated proteins for each metabolite, 225 protein-based prediction models were trained (Supplementary Table 6). We further compared the top 50% high R^2^ metabolite models from gene expression data, and proteomic data, 53.1% (60 of 113) metabolites were detected by both proteomics-based and gene-based models (Fig. 2d,e).

### The regulatory mechanism in metabolic pathways involve more than genes and proteins

As the prior analyses showed, selecting correlated genes effectively improves the prediction performance of models, therefore, we performed similar studies with genes that functioned in metabolism pathways. Metabolism pathway information was obtained from the database SMPDB (https://smpdb.ca/), and the 554 metabolism genes that are in these pathways were selected for further analysis. For every cell line, the GSVA score^30^ was calculated based on the expression of genes and metabolite profiles, respectively, among 32 metabolic pathways that contained at least 5 genes or metabolites. The Pearson correlation scores range from - 0.18 to 0.25 (Supplementary Table 7), suggesting that there is no or only a weak correlation between gene GSVA scores and the metabolite GSVA scores within the same pathway.

Although the pathway level analyses did not reveal a strong correlation between genes and metabolites, we further assessed whether using the 554 genes from SMPDB (Supplementary Table 8) alone can improve prediction. The R^2^ in *LASSO* training models ranged from 0 to 0.69 (Fig. 3a, Supplementary Table 9). While the majority of the metabolism gene-based *LASSO* prediction models showed lower R^2^ than that of global gene-based prediction models, with the median of the training set and testing set was still around 0. We further performed a replication study with 1701 metabolism genes from Reactome database (https://reactome.org/, Supplementary Table 10). However, the performance of prediction models remained similar (Fig. 3b; Supplementary Table 11).

**Figure 3.**
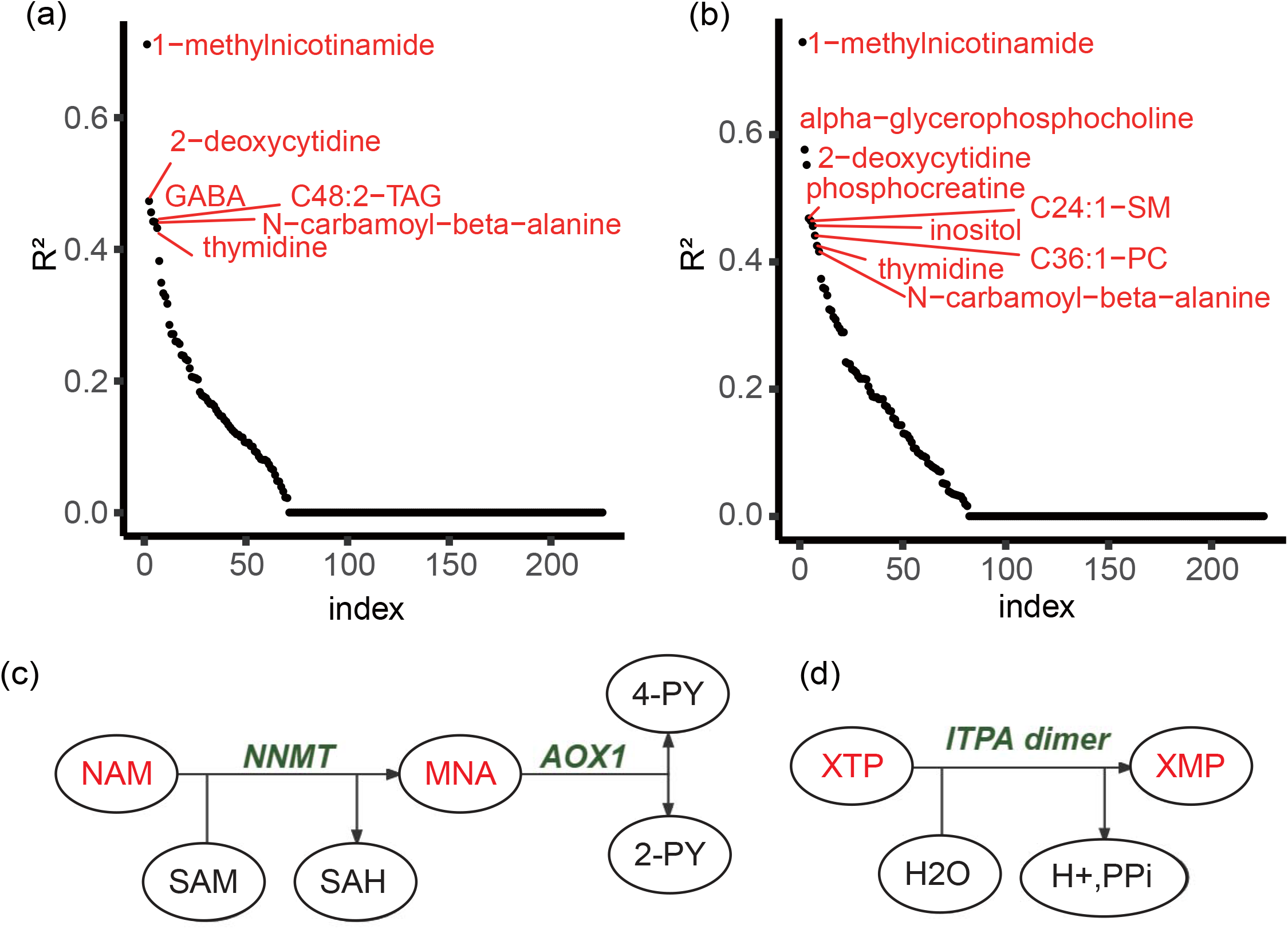
metabolic pathway level analysis result. a) R^2^ distribution of metabolism gene-based LASSO regression prediction models, the metabolism gene list was downloaded from SMPDB and b) Reactome database; c) 1-methylnicotinamide metabolic pathway; d) xanthosine metabolic pathway * MNA: 1-methylnicotinamide; NAM: Nicotinamide; XMP: xanthosine; XTP: xanthine.

As most metabolic networks are complex with multiple enzymes that could result in the catabolism and metabolism of a specific metabolite, we evaluated the importance of also considering quantified metabolite levels within a metabolic pathway. To do so, we generated models that, in addition to gene expression, also included the level of reactant metabolites within the same pathway. Two relatively simple and unidirectional metabolism pathways were chosen as examples. First, we developed a *LASSO* regression model to predict the level of 1-methylnicotinamide (MNA), which is the most well-predicted metabolite, with its upstream reactant Nicotinamide (NAM) and regulated genes (Fig. 3c). The prediction model performance (R^2^) did not change, more importantly, NAM was not selected as a significant predictor. Similarly, we modelled the level of xanthosine (XMP) by the correlated genes and its associated nucleoside xanthine (XTP) (Fig. 3d). Interestingly, XTP became the only valued predictor in the trained model, with the R^2^ increasing from -0.524 to 0.376 in the testing set (Table 2). This example suggests that, in some cases, considering the level of associated metabolites within a pathway is essential for metabolite prediction.

**Table 2.** Prediction power change after adding reactant metabolite level of MNA and XMP metabolism.

| **Prediction Model** | **R<sup>2</sup> in testing set** | **Reactant Coefficient** |
| --- | --- | --- |
| **MNA ~ selected genes** | 0.683 | \ |
| **MNA ~ selected genes + NAM** | 0.683 | \ |
| **XMP ~ selected genes** | -0.422 | \ |
| **XMP ~ selected genes + XTP** | 0.376 | 0.57 |

MNA: 1-methylnicotinamide; NAM: Nicotinamide; XMP: xanthosine; XTP: xanthine.

### Nicotinamide N-methyltransferase is a robust predictor of 1-methylnicotinamide expression only in cancer cells

Among all profiled metabolites, MNA showed the strongest positive correlation with the expression of the gene Nicotinamide N-methyltransferase (*NNMT*) (Supplementary Fig 2B,2C). In both gene-based and protein-based MNA prediction models, *NNMT* was the most effective predictor of MNA level. *NNMT* encodes an enzyme, which methylates nicotinamide to produce MNA^31^.

We tested the MNA prediction model in a breast cancer bulk tissue dataset^25^ with matched gene expression data and metabolomic profiles. To our surprise, though MNA was generally abundant in the cancer samples (Fig. 4a), there was no significant linear relationship between *NNMT* and MNA (Fig. 4b). A recent study deciphered that the gene *NNMT* is mainly expressed in cancer epithelial cells and fibroblasts, while immune cells can take-up MNA released in the microenvironment but seldomly express *NNMT* themselves^32^. As the bulk tissue samples were comprised of different amounts of cancer cells, immune cells and fibroblasts, the positive correlation between MNA level and *NNMT* expression may not be shown in such low cancer purity bulk samples. To test this hypothesis, we used ESTIMATE^33^ to evaluate the purity of those breast cancer samples (Fig. 4c). Bulk tissue samples had relatively low purity (range around 0.3 to 0.9) compared with that of CCLE cell lines (most purity of >0.95). We further compared the relationship between MNA level and *NNMT* expression in the solid cancer cell lines and those from myeloid or lymphoid leukaemia and lymphoma cells. Indeed, the relationship was only present in cancer epithelial cells (Fig. 4d). Generally, solid cancer cells and blood cancer cells show substantially different metabolic profiles (Fig. 4e).

**Figure 4.**
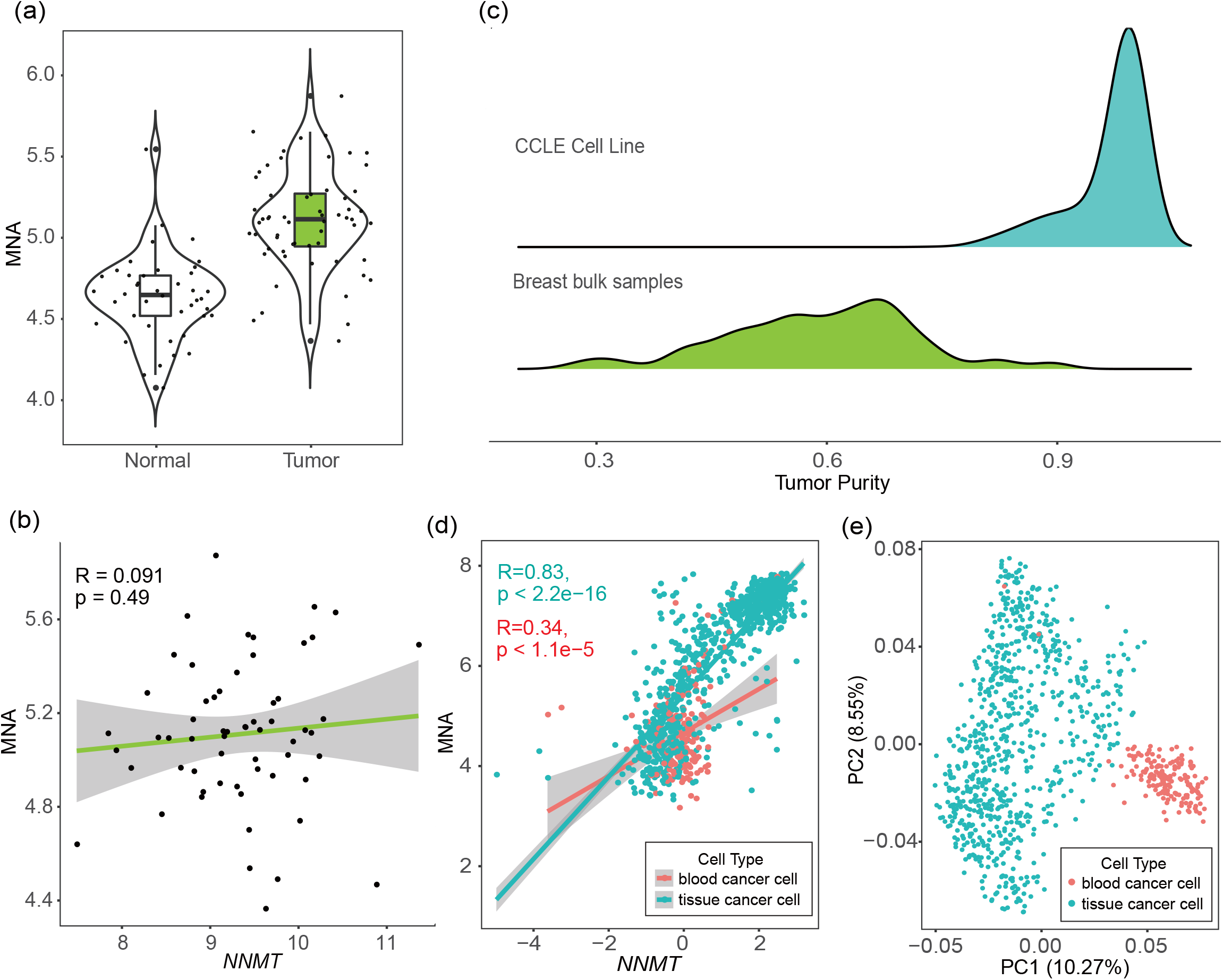
Linear relationship between NNMT and MNA in different dataset. a) MNA level in cancer samples and normal samples from breast cancer cohort (n=102). b) The linear relationship between MNA and NNMT in breast cancer bulk tissue samples. c) cancer purity evaluated by R package ‘estimate’. d) The linear relationship between MNA and NNMT in cancer cell lines. e) PCA plot of CCLE metabolism profiles, solid cancer cells and blood cancer cells show substantially different metabolic profiles.

## Discussion

Previous studies have pointed to the importance of metabolism in cancer development^34^. The initial objective of this study was to clarify whether gene expression can be used to infer metabolic activity and predict metabolite levels. By using the matched gene expression and metabolism profiles from the same cohort, we implemented a pan-cancer and multiple cell-culture media study of 454 human solid cancer cell lines from CCLE. Three approaches to train gene-based models were attempted, however, the metabolism heterogeneity and sample purity enlarge the challenge of metabolite prediction, which was also suggested by previous studies^27,35^. The R^2^ of prediction models largely varies among metabolites, with less than 2% (3 of 225) metabolites that consistently had an R^2^ above 0.5 in both training set and testing set using the optimal modelling method. These results illustrated the unpredictability of most metabolites from gene profiles, even considering prior knowledge such as cell culture conditions and metabolism pathways. Analogous analysis with proteomics profiles resulted in a similar conclusion.

One interesting finding is that MNA maintained good performance in all gene-based and proteomics-based models, under a strict cell type requirement. Meanwhile, MNA was found highly accumulated in cancer cells in a breast cancer bulk tissue dataset, which supports our finding. MNA is predictable by the expression of *NNMT*, probably because of the clear metabolic relationship within the pathway, and the downstream regulated gene *AOX1* is not expressed in most cancer cells^36^. The majority of metabolic activities are not as simple as this, making the prediction of metabolite levels using gene expression challenging. Besides MNA, thymidine and phosphocreatine also have a relatively high R^2^ (>0.5) in the prediction model, while the other 222 metabolites detected in CCLE Cell lines could not be predicted accurately using gene expression data, as they failed to be validated in the testing set. A possible explanation for this might be the complex metabolism activities *in vivo*. Some metabolites are involved in multiple metabolic pathways or in a two-sided metabolism reaction, thus, additional gene sets and side products may be needed to evaluate the reaction rate. Previous studies also point out the bi-directional effect of metabolism and gene expression, making the metabolite level much more unpredictable^15,37^. The interaction between a specific enzyme and regulated gene can not fully explain the metabolic pathway activities, while there are many flexible transcriptional solutions to regulate the metabolism activities and meet the cell survival requirements. Moreover, the enzyme activities could be regulated allosterically and post-translationally, and therefore the gene expression levels of an enzyme do not fully reflect their activities. On the other hand, the metabolism heterogeneity among cells may also influence the prediction result. The metabolism patterns vary among cell types, especially within immune cells^38,39^, the potential interference of cancer purity cannot be ruled out in metabolism prediction in bulk tissue samples.

Our analysis suggests a weak relationship between genes and metabolites. These findings prove that gene expression alone cannot be used to estimate metabolism activity. There are more factors, such as cell types and related metabolites, that should be taken into consideration. This has significant implications on studies that infer metabolism based on gene expression profiling data alone. Nevertheless, a limitation of this study is that the number of metabolites analyzed and the number of cancer samples profiled is still relatively small. Furthermore, the cell line-based data makes these findings less generalizable to bulk tissue samples, although it is likely that predicting metabolite levels from gene expression data will be even more challenging for bulk tissue. Taken together, our study offers some insights into the study of cancer metabolism and highlights the challenges of studying cancer metabolism through gene expression profiling.

## Materials and Methods

### Sample filtering

The profiles of 225 metabolites have been profiled by the CCLE across 928 cell lines^28^, 184 of them were blood cancer cell lines, while the rest were all cancer cell lines from human solid cancer. Only cell lines with adherent growth properties were kept to reduce the potential bias due to cell culture methods. In the end, 454 human solid cancer cell lines across over 24 cancer types with both metabolomic and gene expression profiles were analyzed.

### Gene expression data pre-processing

1,019 Cell lines with transcription RSEM profiles were downloaded from the CCLE database. The transcription features were required to pass the following quality control criteria: 1) only features that can match to Gene symbol were kept; 2) at least 10% cell lines have non-zero values. After filtering, the gene expression data were log10 processed to enlarge the signal. Quantile normalization was performed by R package “preprocessCore” and 13,403 gene features used in the end.

### Estimate coefficient of variation for each metabolite

Coefficient of variation (CV) analysis was based on the computational approach proposed by Reed et al^40^. in 2002, which is defined as the standard deviation (SD) divided by the mean and is used to assay variability. We measured the SD and mean, then calculated CV for each metabolite with R code. The complete CV values are marked in Supplementary Table 1 and Fig 1.

### Train step-wise univariate linear regression models

To illustrate some of the potential considerations for reducing bias and thus optimizing the metabolite prediction performance, we combined 55 main cell culture media components and six clinical traits (inferred ethnicity, patient age, pathology, cancer type, cell lines’ mutation rates, and doubling time(hr)), then constructed both-sided stepwise linear regression models for each metabolite. Before model training, VIF scores were calculated by R package “car” to check the feature multicollinearity.

Those models were trained using the *step* and *lm* functions in R package “stats”, with default setting and direction “both”. Selected features and an R^2^ value were generated for each model automatically. R^2^ represents the coefficient of determination, a value of closer to 1 means better prediction performance of the model. In this step, R^2^ values were used to estimate the component influence.

### Global gene-based LASSO model training

After merging metabolism profiles and gene profiles from the CCLE database, 454 solid cancer human cell lines were analyzed further. This cohort was divided into a training set (n=318) and a testing set (n=136) randomly. We utilized gene expression to predict metabolites in the training set and validated models in the testing set. Univariable linear regression algorithm (R package “glmnet”, lm function) was run for each metabolite for feature selection, the top 10% highly linear-correlated genes were selected as features in a further 10-fold cross validated *LASSO* regression model (R package “glmnet”, cv.glmnet function). R^2^ was used to assess the model accuracy. The result for metabolites prediction models in the training set and testing set are recorded in Supplementary Table 5. In model training, the parameter set as “gaussian” family and alpha as 1, with lambdas tried “10^seq (2, -2, by = -.1)”, features were standardized within the algorithm.

Proteomics-based prediction models were trained using the same pipeline, with proteomics data obtained from a previous study^29^. 351 cell lines had both proteomics profiles and metabolism profiles were further analyzed. We regarded the top 50% high R^2^ metabolite in the testing set as “predictable” metabolites. We found the common “predictable” metabolites from global gene-level prediction models and proteomic-level prediction models by “VennDiagram” R package.

### Estimate Cancer purity

Cancer purity represents the proportion of cancer cells within the cancer tissues. It was estimated from gene expression profiles by R package “estimate” with the default setting.

### GSVA correlation analysis

Gene Set Variation Analysis (GSVA)^30^ is an unsupervised, non-parametric method for predicting variation in gene set enrichment using samples from an expression data set. Based on its statistical principles, we tried to transfer it to metabolism data and measured the metabolic activities in each pathway. A high Pearson correlation rate (correlation rate > 0.9) between metabolism GSVA scores and mean of metabolite level in pathways ensure the feasibility of this analysis. The pathway information was downloaded from SMPDB (https://smpdb.ca/) and the pathways with less than 5 genes or metabolites were filtered with 32 common pathways remaining. GSVA scores based on gene expression and metabolism profiles were calculated by R package “GSVA”, respectively. Therefore, every cell line had one gene GSVA score and one metabolism GSVA score for each pathway. We then used the “cor” function in R to evaluate the correlation between gene GSVA score and metabolism GSVA score. The result is shown in Supplementary Table 7.

### Metabolism gene-based LASSO model training

554 metabolism pathway-related genes were extracted from the SMPDB database (https://smpdb.ca/) and 1701 overlaped metabolism genes were collected from Reactome Database (https://reactome.org/). We trained *LASSO* regression models with those metabolism genes for each metabolite in the training set, and recorded R^2^ distribution from both the training set and testing set. The training was done with the same setting as those in global gene-based *LASSO* regression models.

## Abbreviations

CCLE: Cancer Cell Line Encyclopedia
CPTAC: Clinical Proteomic Tumor Analysis Consortium
GC: Gas Chromatography
LASSO: Least Absolute Shrinkage and Selection Operator
LC: Liquid Chromatography
MS: Mass Spectrometry
NMR: Nuclear Magnetic Resonance
TCGA: The Cancer Genome Atlas

## Acknowledgments

We thank C. Qiao for valuable suggestions on model training.

## Figure legends

**Supplementary Figure 1.**
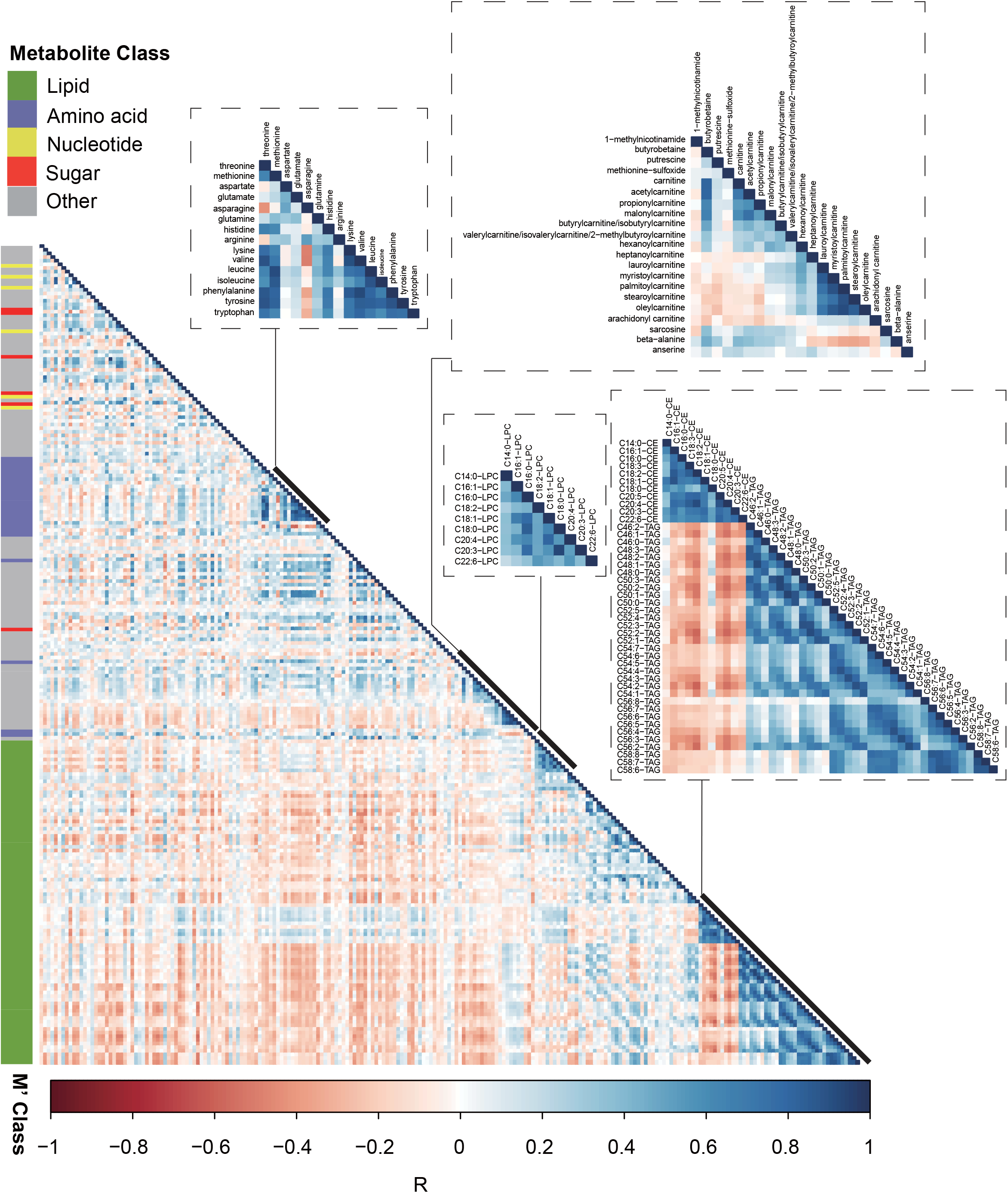
Pearson correlation analyses between every pair of metabolites, R ranges from -0.598 to 1, with mean value of 0.0018; Lipids related to triacylglycerol and ceramide were closer to each other, as well as some amino acids. However, except for some lipids, most of the profiled metabolites show weak correlations (absolute value of Pearson score < 0.5) with other metabolites.

**Supplementary Figure 2.**
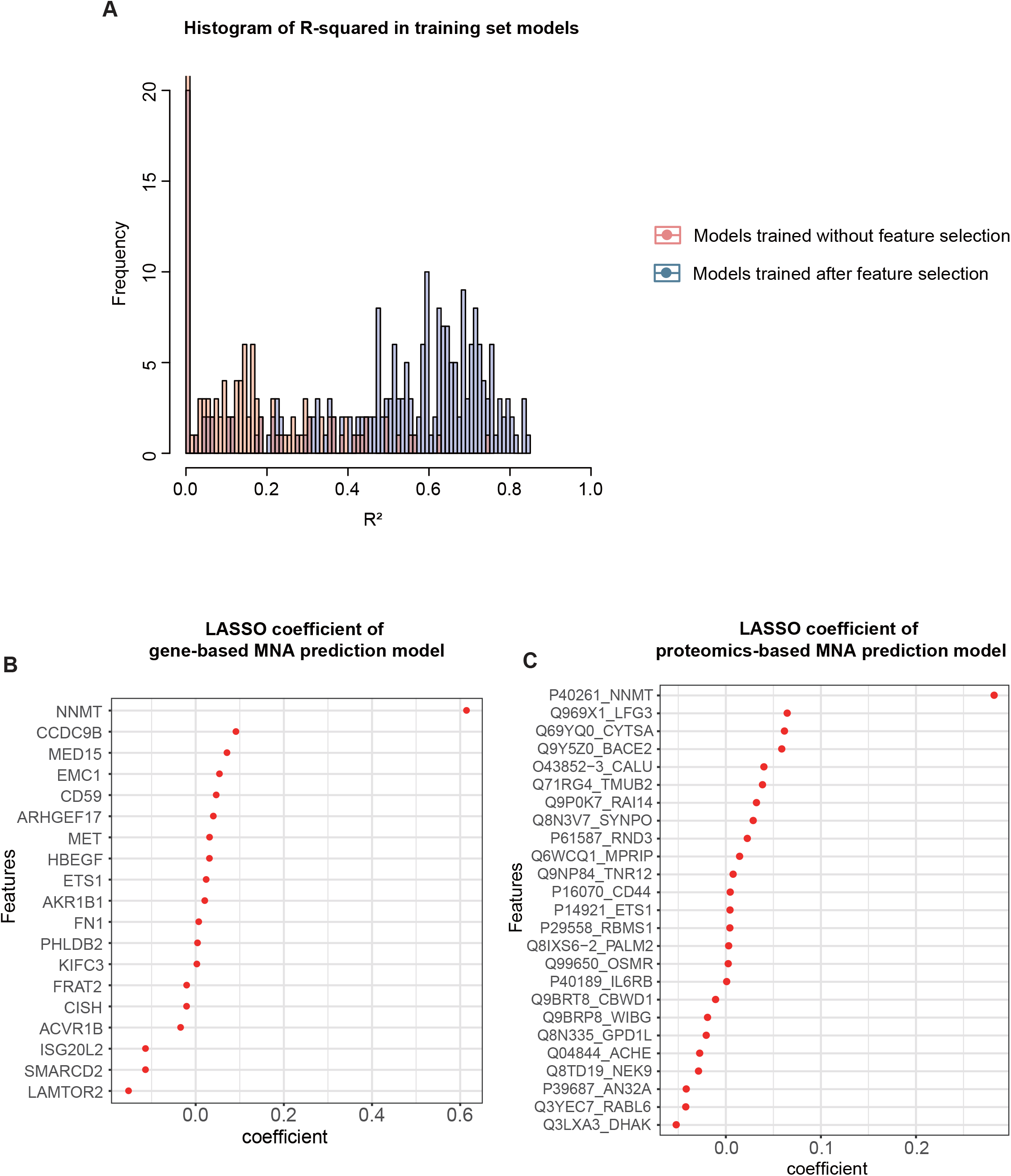
A) histogram of R^2^ from global gene-based multivariable LASSO prediction models with and without feature selection process. B) feature coefficient of NNMT gene and C)NNMT protein.

